# Epistasis in genomic and survival data of cancer patients

**DOI:** 10.1101/130369

**Authors:** Dariusz Matlak, Ewa Szczurek

## Abstract

Cancer aggressiveness and its effect on patient survival depends on mutations in the tumor genome. Epistatic interactions between the mutated genes may guide the choice of anticancer therapy and set predictive factors of its success. Inhibitors targeting synthetic lethal partners of genes mutated in tumors are already utilized for efficient and specific treatment in the clinic. The space of possible epistatic interactions, however, is overwhelming, and computational methods are needed to limit the experimental effort of validating the interactions for therapy and characterizing their biomarkers. Here, we introduce SurvLRT, a statistical likelihood ratio test for identifying epistatic gene pairs and triplets from cancer patient genomic and survival data. Compared to established approaches, SurvLRT performed favorable in predicting known, experimentally verified synthetic lethal partners of *PARP1* from TCGA data. Our approach is the first to test for epistasis between triplets of genes to identify biomarkers of synthetic lethality-based therapy. SurvLRT proved successful in identifying the known gene *TP53BP1* as the biomarker of success of the therapy targeting PARP in *BRCA1* deficient tumors. Search for other biomarkers for the same interaction revealed a region whose deletion was a more significant biomarker than deletion of *TP53BP1*. With the ability to detect not only pairwise but twelve different types of triple epistasis, applicability of SurvLRT goes beyond cancer therapy, to the level of characterization of shapes of fitness landscapes.

**Author Summary:** Genomic alterations in tumors affect the fitness of tumor cells, controlling how well they replicate and survive compared to other cells. The landscape of tumor fitness is shaped by epistasis. Epistasis occurs when the contribution of gene alterations to the total fitness is non-linear. The type of epistatic genetic interactions with great potential for cancer therapy is synthetic lethality. Inhibitors targeting synthetic lethal partners of genes mutated in tumors can selectively kill tumor and not normal cells. Therapy based on synthetic lethality is, however, context dependent, and it is crucial to identify its biomarkers. Unfortunately, the space of possible interactions and their biomarkers is overwhelming for experimental validation. Computational pre-selection methods are required to limit the experimental effort. Here, we introduce a statistical approach called SurvLRT, for the identification of epistatic gene pairs and triplets based on patient genomic and survival data. First, we show that using SurvLRT, we can deliver synthetic lethal interactions of pairs of genes that are specific to cancer. Second, we demonstrate the applicability of SurvLRT to identify biomarkers for synthetic lethality, such as mutational status of other genes that can alleviate the synthetic effect.

## Introduction

Fitness is a measure of replicative and survival success of an individual, relative to competitors in the same population. In this work, we consider the fitness of cells in tumors of cancer patients. Tumors of different patients, also those diagnosed with the same cancer type, display large genomic heterogeneity. Such diverse genotypes of tumor cells result in different tumor fitness, and consequently, different disease aggressiveness and patient survival.

Epistasis is an interaction between genes, and in general refers to departure from independence of effects that their genomic alterations have on a phenotype of interest (12). Bateson (3) first introduced epistasis as a phenomenon of masking of mutation effects. Fisher (16) used the term epistacy for any deviation from additivity in contributions of mutations to the phenotype, where additivity is expected assuming a linear model of genetic alterations as predictors for the phenotype. Beerenwinkel et al. (4) considered epistatic interactions not only among pairs, but also among larger numbers of genes in their contributions to the fitness phenotype. In most general terms, epistasis can be viewed as a property of a mapping from genotypes to their fitness values, called fitness landscape. By estimating epistatic interactions from available data, we can approximate the shapes of fitness landcapes (4).

We now first explain how epistatic interactions are harnessed for the design of efficient anticancer therapy. Second, we propose how the notion of epistasis between triplets of genes relates to therapeutic biomarkers. Modern cancer treatment combines surgery, radiation, chemo-, and also targeted therapy. The advantage of targeted therapy is that it acts against the patient-specific alterations in the tumor. The current state of the art therapies, however, have limited efficacy due to toxicity (18) and rapid development of drug resistance (14, 31). Recently, therapies exploiting synthetic lethal interactions between genes were proposed to overcome these difficulties (2, 8, 20, 23, 32, 33). Synthetic lethality is a negative interaction, where the co-inactivation of two genes results in cellular death, while inactivation of each individual gene is viable. The mechanism behind the success of synthetic lethality-based therapy in cancer is that one gene inactivation already occurs via the endogenous mutation in the tumor cells, and not in the normal cells of the body. Thus, applying a drug that targets the synthetic lethal partner of that gene will selectively kill cancer cells, leaving the rest viable (Fig 1AB). A famous example of clinically exploited synthetic lethal interaction occurs between *BRCA1* and *PARP1*. *BRCA1* mutations disturb error-free homologous recombinational (HR) repair of double strand breaks and the cells become acute sensitive to the lack of single strand DNA break repair, performed by the PARP protein. Thus, in *BRCA1* deficient cells, treatment with a PARP inhibitor is expected to result in high genomic instability and cell death. Indeed, breast and ovarian cancers that harbor *BRCA1* mutations can be treated with drugs targeting PARP1, such as Olaparib (17, 21, 29).

**Figure 1:**
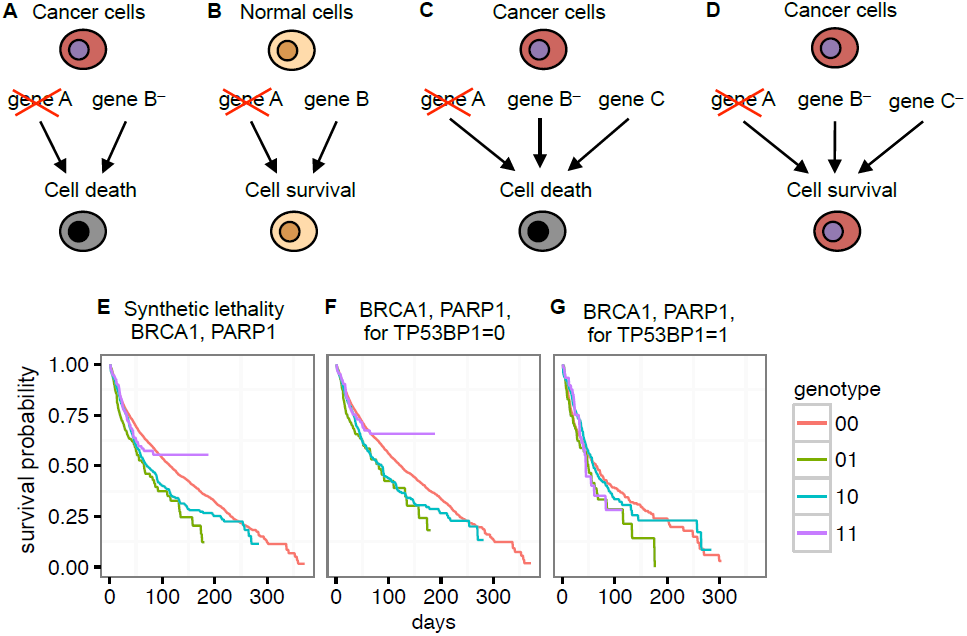
Synthetic lethality and its biomarkers in cancer. **A**,**B** Synthetic lethal partners can be exploited for cancer therapy. In cancer cells (**A**), one partner, here gene B, is already mutated, while in the normal cells (**B**) it is not. Targeting gene A with a drug selectively kills cancer, and not the normal cells. **C**,**D** Mutations in a third gene can serve as therapeutic biomarkers. **C** In the case when the third gene, here C, is not mutated, the therapy targeting gene A in B-mutated cells is successful. **D** Mutation in gene C alleviates the synthetic lethal effect. **E**-**G** Survival functions for patients grouped by the four possible genotypes for **E** the pairwise interaction between *BRCA1*, *PARP1*, **F** the triple epistasis type *a* between *BRCA1*, *PARP1*, conditional on *TP53BP1* not altered, and **G** for the triple epistasis type *b* between *BRCA1*, *PARP1*, conditional on *TP53BP1* alteration. Synthetic lethality is visible in **E** and **F**, but not in **G**.

Synthetic lethality is, however, context dependent. For example, *BRCA1* deficient cell lines were 57 to 133 times more sensitive to PARP1 inhibition, respectively, than normal cells (15). Compared to this dramatic effect, the efficacy of Olaparib therapy on patients was low, since a positive response was observed in less than 50% of BRCA-mutated breast and ovarian cancers (9). This raises the crucial issue of identification of therapeutic biomarkers. For *BRCA1* and *PARP1*, mutation of *TP53BP1* in addition to *BRCA1* was observed to largely restore the function of the HR pathway, and alleviate the synthetic lethal effect (1, 6). Thus, in *TP53BP1* defficient tumors, administrating Olaparib is not justified, and unaltered *TP53BP1* is one of the biomarkers of succes of this therapy. Exactly such dependence of pairwise interaction on the mutational status of a third gene, illustrated in Fig 1CD, is formally represented by so called conditional epistasis, one of the types of triple epistatic interactions studied by Beerenwinkel et al. (4).

Experimental approaches to identification of synthetic lethality in human cancer are overwhelmed by the number of possible interactions (8), which assuming there are 20K genes in the human genome, amounts to c.a. 200 million pairs, and raises to 1.33 × 10^12^ if triplets were considered. High-throughput studies, utilizing short interfering RNA (siRNA) or CRISPR-based screens in human cells, limit the tested pairs to only a subset of all possible (5, 24, 26, 28, 34, 36, 39, 40). The effort and money required for conducting these experiments calls for a pre-selection of synthetic lethal partners for validation based on the computational analysis of existing data. Initial computational approaches were based on the concept of evolutionary conservation and on the knowledge of yeast genetic interactions (11, 13, 27, 41). In our work (37) we aimed at deciphering synthetic sick or lethal interactions from somatic alteration, expression and survival data of cancer patients. To this end, we identified such gene pairs whose aberrations or expression levels were mutually exclusive, i.e., their simultaneous occurrence was under-represented in the tumor cells, given their individual prevalence. Additionally, we checked whether their simultaneous inactivation coincides with increased patient survival. Jerby-Arnon et al. (22) combined different predictors of synthetic lethality, two of which, referred to survival of the fittest (SoF) and coexpression, were most successful. Following the mutual exclusivity principle, SoF identifies synthetic lethal gene pairs when their co-inactivation occurs significantly less frequently than expected. Coexpressed genes usually participate in closely related biological processes, which should be the case for synthetic lethal partners.

Here, we introduce SurvLRT, an approach for identification of epistatic gene pairs and triplets in human cancer. We propose a statistical model based on Lehman alternatives (25), which allows to estimate fitness of tumors with a given genotype from survival of carrier patients. We assume that a decrease of fitness of tumors due to a particular genotype is exhibited by a proportional increase of survival of the patients.Accordingly, for synthetic lethal genes, such as *BRCA1* and *PARP1* (Fig 1E), the survival of patients with the double mutation should be longer than expected from survival of patients with only single mutations and of patients without mutations of those genes. Based on these assumptions, we introduce a likelihood ratio test for the significance of a given pairwise or triple epistatic interaction. In the test, the null model assumes that there is no epistasis and the gene alterations are independent, while the alternative assumes otherwise. The approach can detect both positive and negative interactions. Compared to our previous method (37), SurvLRT offers a more natural interpretation of the notion of fitness, as well as a direct statistical test for the significance of epistasis. We provide the theory for a total of 13 different epistasis types on pairs and triplets of genes defined by Beerenwinkel et al. (4), and illustrate testing and interpretations for three of them on patient data. First, we analyze the sensitivity and power of SurvLRT for all considered epistasis types in a controlled setting of simulated data. Next, we show that, compared to SoF and coexpression, our method performs favorably in predicting known pairwise synthetic lethal interactions. Finally, we apply SurvLRT to detect therapeutic biomarkers, first by recapitulating *TP53BP1*, the known biomarker for therapies based on the *BRCA1*, *PARP1* interaction (Fig 1FG), and second by identifying a genomic region deleted in tumors as a new and even more significant biomarker than *TP53BP1*.

**Table 1:** Conditions for the considered types of pairwise and triple epistatic interactions.

| Type | Condition | Description |
| --- | --- | --- |
| pairwise | $\Delta_{01} \cdot \Delta_{10} \neq \Delta_{00} \cdot \Delta_{11}$ | – |
| *a* | $\Delta_{010} \cdot \Delta_{100} \neq \Delta_{000} \cdot \Delta_{110}$ | conditional |
| *b* | $\Delta_{011} \cdot \Delta_{101} \neq \Delta_{001} \cdot \Delta_{111}$ | conditional |
| *c* | $\Delta_{001} \cdot \Delta_{100} \neq \Delta_{000} \cdot \Delta_{101}$ | conditional |
| *d* | $\Delta_{011} \cdot \Delta_{110} \neq \Delta_{010} \cdot \Delta_{111}$ | conditional |
| *e* | $\Delta_{001} \cdot \Delta_{010} \neq \Delta_{000} \cdot \Delta_{011}$ | conditional |
| *f* | $\Delta_{101} \cdot \Delta_{110} \neq \Delta_{100} \cdot \Delta_{111}$ | conditional |
| *g* | $\Delta_{011} \cdot \Delta_{100} \neq \Delta_{000} \cdot \Delta_{111}$ | marginal |
| *h* | $\Delta_{010} \cdot \Delta_{101} \neq \Delta_{001} \cdot \Delta_{110}$ | marginal |
| *i* | $\Delta_{010} \cdot \Delta_{101} \neq \Delta_{000} \cdot \Delta_{111}$ | marginal |
| *j* | $\Delta_{011} \cdot \Delta_{100} \neq \Delta_{001} \cdot \Delta_{110}$ | marginal |
| *k* | $\Delta_{001} \cdot \Delta_{110} \neq \Delta_{000} \cdot \Delta_{111}$ | marginal |
| *l* | $\Delta_{011} \cdot \Delta_{100} \neq \Delta_{010} \cdot \Delta_{101}$ | marginal |

## Materials and methods

### Types of epistatic interactions

Given *n* genes, their genotype is a tuple *g ∈* {0, 1}^*n*^. Here, we assume that *g*(*i*) = 1 if and only if gene *i* acquired a somatic deletion in the tumor genome. For a pair of genes, their possible genotypes are, in lexicographical order, 00, 01, 10, 11. Denote the fitness of genotype *g* by Δ_*g*_ and its logarithm by *δ*_*g*_. Assuming no interaction between a pair of genes, we expect no deviation from additivity in log fitness, i.e., that *δ* = *δ*_00_ *-δ*_01_ *-δ*_10_ + *δ*_11_ = 0 (4). Equivalently, epistasis between a pair of genes occurs when Δ_01_ Δ_10_ ≠ Δ_00_ Δ_11_ (Tab 1). In the following, we call the deviation *δ* the epistatic effect size. *δ >* 0 indicates a positive, while *δ* < 0 indicates a negative epistatic interaction. The above epistasis definition can be extended from pairs of genes to triplets. Beerenwinkel et al. (4) list twenty types of possible three gene interactions, denoted *a-t*. Conditional epistasis (*a-f*) occurs among a pair of genes, conditioned on the fact that another gene is fixed (Tab 1). For example, conditional epistasis *a* is an interaction between the first and the second gene in the triple, given that the third gene is not mutated. Conditional epistasis of type *b* is defined analogously, but conditioning on the fact that the third gene is mutated. The epistasis types *g − l* are called marginal epistases (Tab 1). For example, marginal epistasis *k* corresponds to the synthetic lethal effect between one mutation in the third gene in the triplet, and two mutations: in the first and in the second gene. In summary, the epistasis types discussed here (listed in Tab 1) are defined by four genotypes *g*_0_ *< g*_1_ *< g*_2_ *< g*_3_ (in lexicographical order), which satisfy the equation *g*_0_ *-g*_1_ *-g*_2_ + *g*_3_ = 0 (the zero vector), and by the epistasis condition Δ_*g*_1 *·* Δ_*g*_2 *≠* Δ_*g*_0 *·* Δ_*g*_3. In this work, we ignore the epistasis types *m - t*, as their defining expressions can be derived as sums over expressions for the types *a - l*, e.g.,*m*: = -(*a* + *k*) (4).

### SurvLRT

We now introduce a survival model of tumor fitness, based on the concept of Lehman’s alternatives (25). We assume we are given survival times of patients and we record the genomic alterations of genes in their tumors. Let *T* be a random variable indicating survival time of a single patient. We assume that the distribution of *T* for healthy individuals has cdf *F* and survival function *G*(*t*) = 1 *F -* (*t*) = P(*T > t*). Our model assumes that the survival function for patients with tumor genotype *g* reads

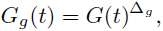

where Δ_*g*_ *>* 0 corresponds to the fitness of the tumor cells for that patient. Note, that since *G*_*g*_(*t*) *∈* [0, 1], the larger the exponent Δ_*g*_, the lower the *G*_*g*_(*t*) becomes. This is in agreement with the assumption that with increased tumor fitness, the disease becomes more aggressive, and patient survival decreases accordingly, as compared to the reference survival. Importantly, accumulation of mutations of independent (not interacting) genes, should correspond to raising the initial survival function *G*(*t*) to consecutive exponents. For example, for a pair of such not interacting genes, we expect that 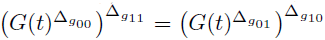, which is of course equivalent to Δ_01_ *·* Δ_10_ = Δ_00_ *·* Δ_11_. In general, the condition for an epistatic interaction of any type (Tab 1) with given genotypes *g*_0_ *< g*_1_ *< g*_2_ *< g*_3_ is satisfied when

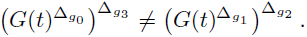

The model presented above allows us to develop a surprisingly simple method, which we call SurvLRT, for evaluating epistasis directly from cancer patient data. Indeed, suppose that 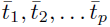 are survival times for patients with genotypes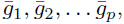 respectively. Let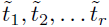 be times of last contact with patients with genotypes 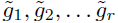, respectively. This part of data is censored and we do not know the exact survival time of the *j*^*th*^ patient - we only know that it is longer than 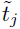. In the following, to simplify the notation,we use Δ_*i*_ to shortly denote Δ_*gi*_. To test for a selected epistasis type, either pairwise, or one of the triple epistasis types over genotypes *g*_0_ *< g*_1_ *< g*_2_ *< g*_3_, we follow two steps. First, we compute the likelihood ratio for the null hypothesis Δ_0_ *·* Δ_3_ = Δ_1_ *·* Δ_2_ and the alternative Δ_0_ *·* Δ_3_ ≠ Δ_1_ *·* Δ_2_. Next, we obtain the maximum likelihood estimators of Δ_0_, Δ_1_, Δ_2_, Δ_3_, determining whether the interaction is positive (*δ >* 0) or negative (*δ <* 0). The likelihood ratio equals

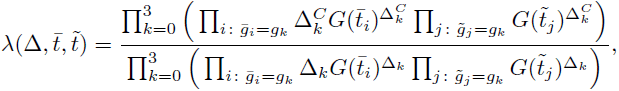

where 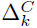 denote the parameters of the null model, which are constrained to satisfy 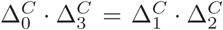 (Appendix). Under the technical assumption that *F* has a density, the maximum likelihood estimators are given by the formula

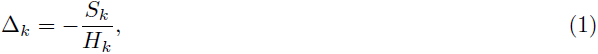

where 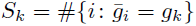 and 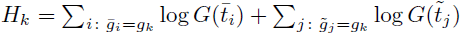 (Appendix). The constrained parameters in the null model 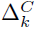 are given by the following formulae for 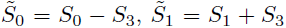 and 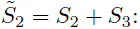

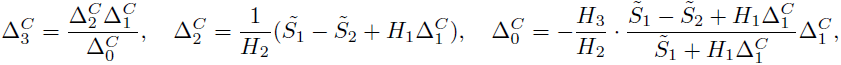

where 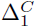 is a solution of the quadratic equation

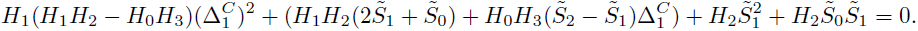

This equation, in combination with the formulae for 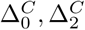 and 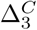, may return two solutions for parameter values, and we chose the one in (0, ∞)^4^. If both do, we choose the one for which the value of the log likelihood is greater (Appendix).

In the case when the data contains only uncensored cases, by the Wilks’ theorem, we can safely assume that, asymptotically,

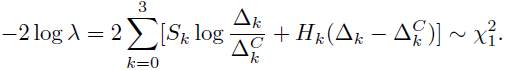

Addition of censored cases does not cause large deviations from this assumption (Fig S1). This allows computing p-values and confidence intervals for the tested epistatic interactions.

### Data processing

The data for 9899 patients of 32 cancer types generated by the Cancer Genome Atlas (TCGA) (38) was downloaded using the cgdsr R package from the cBioPortal (7). The data were organized into a patient per gene binary matrix, where the patient acquired value 1 for the gene if the gene was deleted in that patient’s tumor, and 0 otherwise. Deletion was called for a gene for which GISTIC (30) returned value -1 or -2, and its mRNA level was concordant with its alteration, i.e., the median mRNA expression of that gene was smaller for patients with deletion than in the whole population, as assessed with a Wilcoxon test (lower tail p-value ≤ 0.05). After filtering out patients and genes with excess of missing values, we analyzed a dataset covering 9484 patients of 27 cancer types (Tab S1). A smaller cohort of 2942 breast cancer patients (Tab S2) was used to validate SurvLRT predictions against a set of ground truth interactions tested experimentally in breast-cancer cell lines (Tab S3). In all analyses below, we approximate the reference survival function *G*(*t*) using Kaplan Meier estimate of the survival function for a cohort of 582105 patients that died of other reasons than cancer, collected by the Surveillance, Epidemiology, and End Results Program (35) (Tab S4).

### Controlling false positive rate

When applied to make new epistasis predictions, SurvLRT needs to test all possible interactions between all possible pairs or all possible triplets of genes in question, which raises the problem of multiple hypothesis testing. Moreover, the patient genotypes are defined based on copy number alterations, and several genes may be deleted together within one genomic region. Thus, false positive interactions for genes that happen to be within the same genomic region as the truly interacting gene may be identified. Finally, false positives may occur in the case when the epistasis interaction is tested for a cohort of mutliple cancer types. Each cancer type has its characteristic surivial times, and a cancer type bias for our test may occurr when a cancer type with particularily long or short survival dominates any of the patient groups corresponding to the four genotypes *g*_0_, *g*_1_, *g*_2_ and *g*_3_ considered in the test, since the test statistic may artificially be drawn towards signifficant values. Thus, to control for the false positive rate we take three measures. First, we apply Benjamini-Hochberg correction to the SurvLRT p-values for each tested gene interaction. Second, we group the interactions with the same p-values together so that they are defined per region, containing several genes commonly deleted in patients, and not per individual genes. Third, we control for the cancer type bias. Specifically, denote *w*_0_, *w*_1_, *w*_2_, *w*_3_ the proportions of the respective genotype carriers to the patient total, with Σ*g w*_*g*_ = 1, and *v*_1_, *v*_2_, *…, v*_*m*_ the proportions of cancer types to the patient total, where Σ*c v*_*c*_ = 1 and *m* is the number of cancer types. Under the assumption that the cancers are distributed evenly across the genotypes, we expect the proportion *q*_*gc*_ of patients with genotype *g* and cancer type *c* to satisfy *q*_*gc*_ = *w*_*g*_ *· v*_*c*_ for each *g ∈ {*0, 1, 2, 3*}* and *g ∈ {*1, *…, m}*. To check whether the cancer types in our cohort are evenly distributed across the genotype carriers, we compare the expected proportions *q*_*gc*_ to the proportions 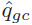 observed in the data. This cancer bias test is conveniently visualized with the expected and the observed proportions on the plot coordinates, and the respective points following the *y* = *x* line if the bias is avoided and the distribution of cancer types across the genotypes is even.

## Results

### Simulations

To evaluate the performance of SurvLRT in a controlled setting, we simulated survival data and fitness values for all types of epistasis among triplets of genes. We first assessed the accuracy of parameter estimation (Fig 2A) for different sample sizes *s ∈*{3, 10, 50, 100, 300, 1*K,* 3*K,* 10*K*} and for two different fractions of censored cases in the data (33% or 66%; the second percentage is more realistic: in the TCGA pan-cancer data there are 63.75% censored cases, and 56.5% in the breast cancer cohort). Here, we investigated whether the parameters obtained using formula (1) agree with the values fixed in the simulations. It was thus enough to simulate one group of patients of a given sample size *s*, assuming they share the same genotype. To this end, we generated *s* observations of survival times *T* at random from the distribution *G*(*t*)^Δ^, where Δ = 1. Next, the censoring times *Y* were sampled from a truncated (up to 40 years) exponential distribution, *Y* ∼ Exp(*c*), where the parameter *c* was set so that the mean percentage of censored observations was either 33% or 66%. In the case when *Y < T*, the last time of follow up for that patient was fixed to *Y*, and the patient was flagged as censored. Otherwise, the patient was flagged as dead, and the time to death was fixed to *T*.

**Figure 2.**
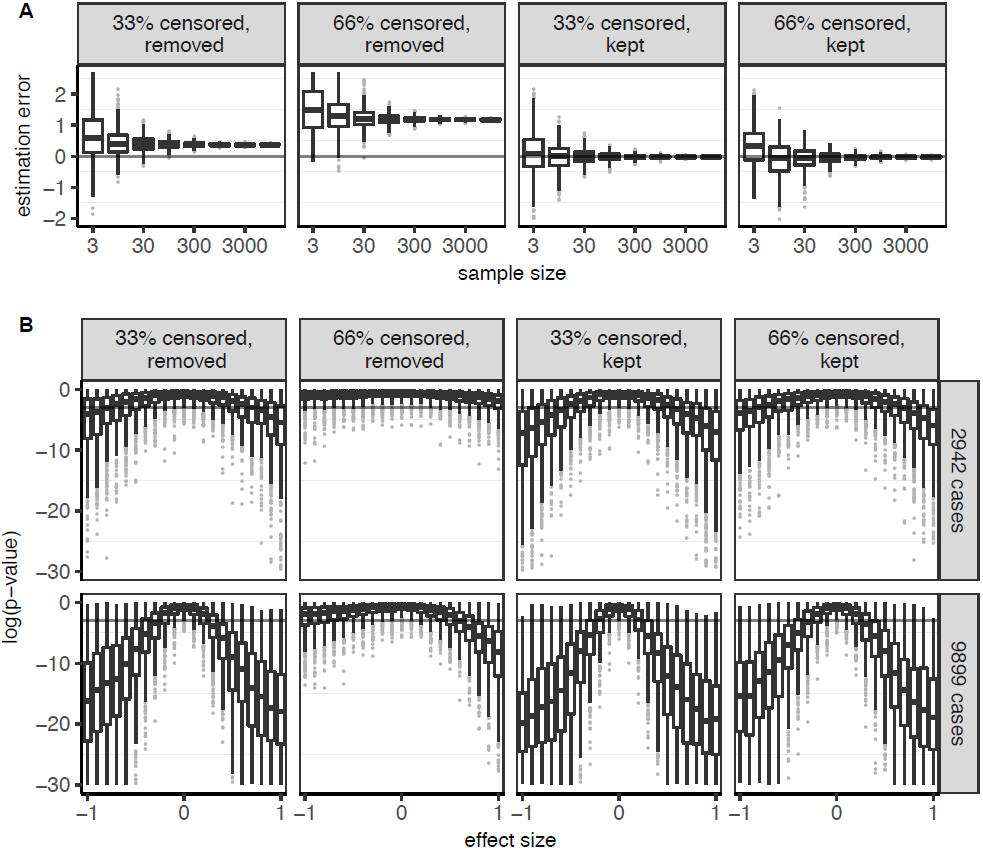
SurvLRT performance on simulated data. **A** Box plots showing 25th, 50th and 75th percentiles (horizontal bars), and 1.5 interquartile ranges (vertical line ends) of log true to estimated fitness ratios (y-axis) as a function of sample sizes (x-axis), in the case when 33% and 66% of patients are censored and removed from the samples (first and second column, respectively) and when 33% and 66% are censored and kept (third and fourth column). Gray lines at 0 mark the level where the estimated equal the true parameter values. **B** Box plots of SurvLRT p-values as a function of epistatic effect size, for two different patient cohort sizes (rows). Gray lines mark the p-value 0.05. Columns as in **A**.

One of the important aspects of survival data analysis is whether to take the censored data into account. Removing the censored cases from the sample and ignoring them in our model lowers the sample size, decreasing the power of the test, and introduces bias in parameter estimation. This bias is more profound, when the percentage of censored cases is higher (Fig 2A, first two columns). The removal of the censored cases changes the distribution of the data: we obtain survival times not from the distribution of *T* itself, but from the distribution of *T* conditioned on the event *T < Y*. Keeping the censored cases maintains the power of the test, and allows accurate parameter estimation (Fig 2A, last two columns). Regardless of the percentage of censored cases, when they are kept in the sample, the median log ratio of estimated to simulated parameter values is 0. The variance of this log ratio decreases substantially as the size of the sample increases, and while some of the parameter estimates from only three samples are unreliable, already for 300 samples all estimates are very close to their true values.

Next, we analyzed SurvLRT p-values as a function of effect size, again for different percentages of censored cases, and for two different total sizes of simulated patient cohorts (Fig 2B). In this analysis, for each simulated triple epistasis type, we fixed the parameters Δ_0_ = 1, Δ_1_ = *e*^0.2^, Δ_2_ = *e*^0.3^, and we set Δ_3_ such that log(Δ_3_) = log(Δ_1_) + log(Δ_2_) + *δ*, where *δ* was the value of the tested effect size. For each gene in the triple, we sampled *k*, the number of patients which had this gene mutated, according to the mutation frequencies observed in the real cancer patient cohort. Next, we chose *k* patients at random to have this gene mutated, fixing their value for this gene to 1, and for all remaining patients to 0. As a result, we obtained patient genotypes for the simulated triple. For each patient with genotype *g*_*i*_, we sampled survival time *T* from the distribution 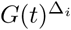, together with a random variable Y ˜ Exp(*c*), with *c* fixed to obtain the wished percentages of censored patients (33% or 66%). As above, the resulting observation for the patient was taken as the min(*T, Y*), and the patient was flagged as censored if *Y < T*, and as dead otherwise. The simulation was repeated first assuming the total number of patients was 2942 (Fig 2B first row), equal to the number of patients in the breast cancer cohort analyzed below, and second for the total number of 9899 patients from the TCGA (second row).

With removal of censored cases from the data it becomes increasingly difficult for SurvLRT to detect epistasis, even for large effect sizes (Fig 2B first two columns). When the censored cases are kept, the power of the test is greatly improved (Fig 2B last two columns). In this case, an increase in the percentage of censored cases from one third to two thirds results in a slight decrease of the power, but by far less dramatically than their complete removal. For the smaller number of around 3K patients, the power of the test is generally low, and increases substantially when a larger patient cohort of around 10K is analyzed. In all scenarios, regardless of the cohort size and share of censored cases, the test correctly returns the largest p-values and does not call epistasis when the effect sizes are 0. In summary, the simulations indicate that the censored cases should be taken into account in the SurvLRT model, to gain advantage of larger sample sizes and to ensure correct parameter estimation. Moreover, for the currently available sizes of single cancer cohorts, like the breast cancer, SurvLRT will return significant p-values only for large epistatic effects, and cohorts as large as the pan-cancer are required to increase the power.

In this and later sections, the reference survival function *G*(*t*) is estimated from patient data (Methods). To assess how this survival function affects the reported results, we estimated an alternative *G*(*t*) from survival times sampled from the exponential distribution, keeping its mean equal to mean survival in the patient data. Both the error of parameter estimates and the power of the tests do not depend on which of the alternative forms of the survival function *G*(*t*) we used, with the only exeption that the spread of the outliers was larger, when the survival times were sampled from the exponential distribution (Fig. S2).

### SurvLRT predictions from patient data agree with experimental results on cell lines

To demonstrate the predictive power of SurvLRT on an independent dataset, we tested its predictions from patient data against a set of gene pairs, whose synthetic lethal interaction was previously investigated using siRNA screens on cancer cell lines. The set comprised 963 pairs, consisting of *PARP1* and its partner genes studied by either Lord et al. (26) or by Turner et al. (39), which we were able to map to a unique official symbol, defined by the Human Genome Organisation (HUGO) Gene Nomenclature Committee (19) (Tab S3). The pairs were called synthetic lethal if a) the partner gene, such as *BRACA1/2* or *ATM*, was mentioned by (26) or (39) as previously reported synthetic lethal with *PARP1*, or b) when targeting the partner gene with two or more different siRNAs sensitized to KU0058948, the PARP1 inhibitor utilized in both studies. Otherwise, the gene pairs were flagged as noninteracting. Since the siRNA experiments were conducted on breast cancer cell lines, we applied SurvLRT to the breast cancer cohort (Tab S2). If for a given gene pair SurvLRT identified a negative interaction, we assigned the pair a score equal to the test statistic *λ*. Otherwise, we assigned it a score - *λ*, and we ranked the pairs in the decreasing order of their scores. In this way, higher score indicated more evidence for synthetic lethality. For comparison, on the same dataset, we ranked the genes by scores from three previous methods: 1) by the decreasing absolute coexpression of genes in the pairs, 2) by the statistic of a Wilcoxon test used to assess whether the co-inactivation genes in the pairs occurs significantly less frequently than expected (SoF; (22)), and 3) by the previously introduced S-scores, tailored for predicting such siRNA-based experiments ((37); Appendix). The predictive power on the experimentally verified gene pairs was assessed with the area under the receiver operating characteristic curves (Fig 3A). The very simple predictor based on coexpression achieved surprisingly good results (AUC 0.63). Still, SurvLRT, with AUC of 0.695, outperformed the S-score (AUC 0.6), as well as coexpression and SoF (AUC 0.59) in predicting synthetic lethality. We note, however, that although SurvLRT obtains overall higher AUC, for false positive rate smaller than 0.25 coexpression or SoF give higher true positive rate than SurvLRT. In addition, we also checked the performance of SurvLRT without the concordance check of per gene deletion calls with gene expression data (Methods). Without the check the AUC for SurvLRT decreased, but by less than 0.01. Although overall the compared AUCs are just moderate, reflecting the limits in power of our and other approaches, these results indicate that there is detectable signal of synthetic lethality in patient data.

**Figure 3.**
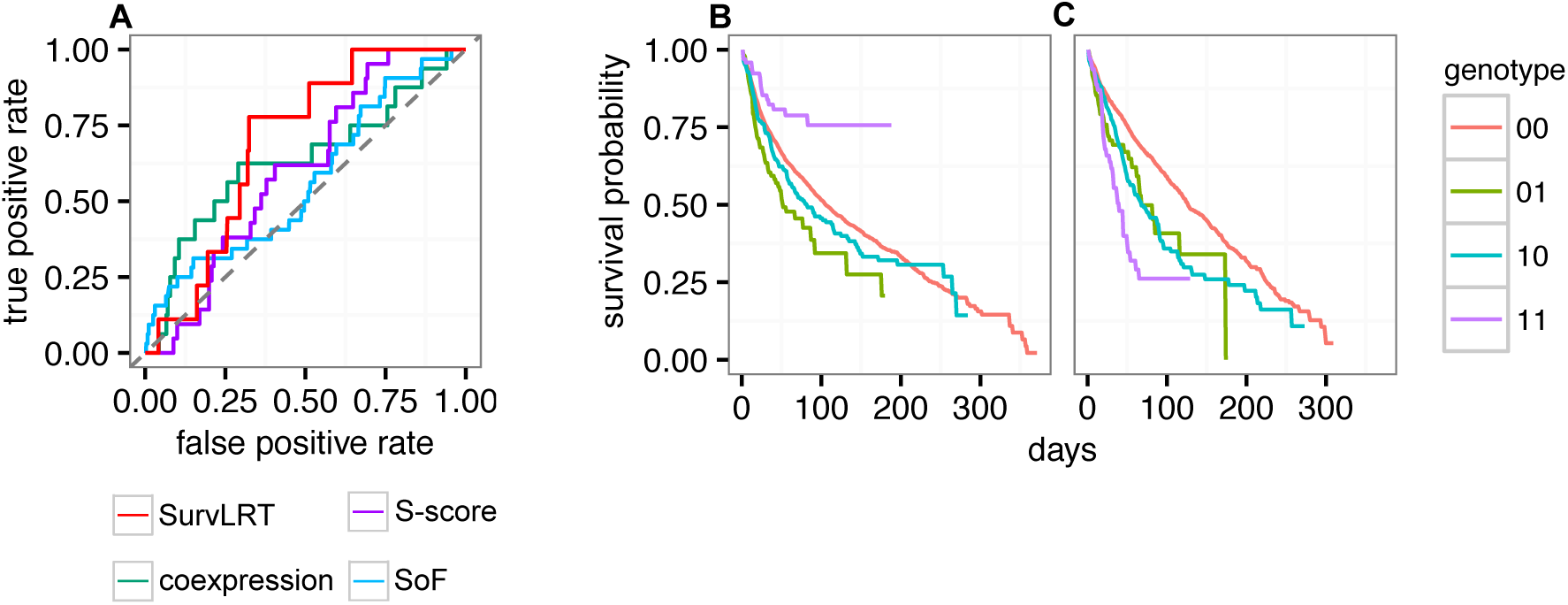
SurvLRT results on patient data. **A** Predictive performance of SurvLRT (red), coexpression (green), SoF (blue) and a random classifier (gray dashed) on experimentally verified synthetic lethal interactions. **B**, **C** Survival functions for patients grouped by the four possible genotypes of the pair *BRCA1*, *PARP1*, and having the newly identified biomarker region not altered (**B**), or having this region deleted (**C**). Synthetic lethality is visible in **B** but not in **C**.

### Using SurvLRT on triplets of genes for biomarker identification

Finally, we investigated whether biomarkers for synthetic lethal interactions can be identified from patient survival data using SurvLRT. First, we applied SurvLRT to test the masking of synthetic lethality between *BRCA1* and *PARP1* by *TP53BP1* alteration (1, 6). To this end, we tested conditional epistasis types *a* and *b* for these three genes on the pan cancer cohort (Tab S1), where the sample size was large enough to obtain a satisfactory power of the test. Recall that type *a* represents epistasis between the first two genes, here *BRCA1* and *PARP1*, conditional on the lack of mutation in the third gene, here *TP53BP1*. As expected in this case, when testing for type *a*, SurvLRT called significant (p-value 0.0003) and negative conditional epistasis (Fig 1F). Additional check assured that the significance of the test is not due to a cancer type bias (Fig. S3). Type *b* represents epistasis between the first two genes, but conditional on the presence of mutation in the third gene. In the test for type *b*, SurvLRT correctly identified that the synthetic lethality interaction is no longer present (Fig 1G). In fact, for these three genes, conditional epistasis of type *b* is positive, although with an insignificant p-value of 0.97. This test showed, but only small, violations to even cancer type distribution (Fig. S3). Thus, SurvLRT in both tests returned results in accordance with biological knowledge. It is worth noting that in this analysis we are unable to compare to any existing approach, since there are no other methods that explicitly test for epitasis between triplets of genes.

Second, to make new predictions, we determined whether any of 1856 pan-cancer genes, significantly and concordantly deleted in tumors (10), could also play a role of biomarkers for the pair *BRCA1*, *PARP1*. To this end, for each gene *G* we applied SurvLRT to test the conditional epistases of type *a* and type *b* for the triple *BRCA1*, *PARP1*, *G*. As potential biomarkers we considered, as in the case of *TP53BP1*, such *G* for which type *a* was significant negative and type *b* was insignificant, after correction for false discoveries (Methods). This procedure identified not a single, but ten genes, *CBFB*, *ZFHX3*, *MLKL*, *CSNK2A2*, *CTCF*, *CDH1*, *FUK*, *TK2*, *PSKH1*, *WWOX*, which are deleted together within one region, as the most significant biomarker (negative conditional epistasis type *a* Benjamini-Hochberg adjusted p-value 6.35e-5, lower compared to the adjusted 0.002 p-value when *TP53BP1* was tested). The conditional epistasis type *b* turned out to be positive, but with a high adjusted p-value of 0.87. Both tests for the epistasis types *a* and *b* were clearly free of cancer type bias (Fig. S3). Thus, according to patient survival data, the deletion of the region harboring these genes could be the determinant of the success of the therapy using PARP inhibitors on *BRCA1* deficient tumors. Indeed, patients with double *BRCA1*, *PARP1* inactivation survive longer than expected when this region is not altered (Fig 3B), and they do not when the region is deleted (Fig 3C). All tested interactions with the p-value smaller than the p-value for *TP53BP1* as the third partner, are listed in Tab S5. The runtime of both the tests of two epistasis types for 1856 gene tiplets, described in this section, as well as the tests for experimentally verified gene pairs, described in the previous section, was less than minutes on a 8GB RAM laptop with a dual core processor.

## Discussion and conclusions

This paper presents SurvLRT, a statistical approach to resolving epistasis from genomic and survival data of cancer patients. SurvLRT has several important benefits. Modeling survival functions using Lehmann alternatives allowed for a natural interpretation of the model parameters as tumor fitness values. Based on this model, we introduced a likelihood ratio test for epistasis that directly tests the log linearity in log fitness of gene mutations, expected when there is no interaction present. With a unified approach it can not only test for epistasis between pairs, but also for interactions among triplets of genes. It detects whether the interaction is positive or negative. Apart from direct estimation of tumor fitness values, it assesses the epistatic effect size, and returns p-values for the significance of the tested epistatic interaction. The advantage of our analysis of cancer patient data over studies performed on cell lines is that we gain access to the more realistic context, where fitness of tumor genotypes depends on their real advantage gained in their natural environment, and is expressed in patient survival.

For cancer, assessment of epistasis has crucial therapeutic implications. Pairwise synthetic lethal interactions are already successfully exploited in the clinic, and our analysis showed the utility of SurvLRT in mining survival data for evidence of synthetic lethality. In addition, we introduced the concept that biomarkers for synthetic lethality-based therapy can formally be defined as conditional epistasis between triplets of genes, and we showed that SurvLRT can correctly identify such epistasis in the data.

Markedly, the utility of SurvLRT does not limit exclusively to these two cancer applications. In its full functionality, SurvLRT evaluates both pairwise and twelve different types of triple gene epistasis. Thus, in general, our approach can be utilized to approximate the shapes of fitness landscapes, which are determined by the epistatic interactions (4).

It is important to note, however, that by its nature survival data of patients does not provide evidence for all existing epistatic interactions. In particular, if deletions of strictly synthetic lethal genes would co-occur in cancer cells, these cells should disappear from the tumor. Therefore, the survival of patients with the double mutant genotypes would not be available for assessment. Indeed, some of the known synthetic lethal pairs could not be analyzed using SurvLRT, as the required genotypes were not present in the cohort. In general, the fact that we can only access the data of surviving tumors implies, that we can only detect a relatively mild signal of whether the co-occurrence of mutations results in unexpectedly decreased or increased tumor fitness. Thus, compared to the studies on cell lines, analysis of tumor data, although more realistic, may allow less sensitive detection of negative epistasis. Still, on those gene pairs where the genotype data was available, SurvLRT proved to correctly predict synthetic lethality. On top of that, the subtle signals in survival data correspond to small epistatic effect sizes. Our simulations show, that statistically, such small effects can better be picked up when the analyzed cohort is larger. Given the research activity in this area, the collection of cohorts will continue to grow.

Taken together, this contribution makes an important step forward in computational prediction of epistatic interactions. In our mind, predictions of SurvLRT and other approaches alike are meant to eventually guide the experimental effort in browsing the immense space of possible interactions to validate. Our results show that SurvLRT is able to find evidence for epistasis in cancer survival data and thus pinpoint the plausible hypothesis to test.

## Appendix Formulae for the likelihood ratio and parameter estimators

Assume the settings from the *Methods* section in the main text. Recall that the random variable *T* denoting a healthy individual’s lifetime has survival function *G*(*t*), and assume it has a density *f*. With the model assumption that for patients with tumor genotype *g* the survival function is given by 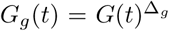, the corresponding distribution function *F*_*g*_ reads

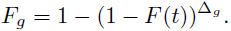

Thus, the density is given by

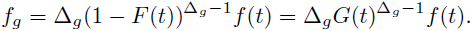

Therefore, the likelihood of the survival data 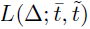, for the uncensored and censored cases, denoted 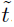 and 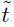, respectively, is given by the formula

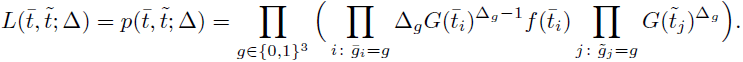

Its logarithm is equal to

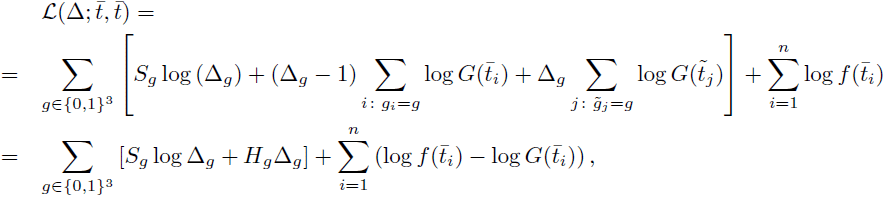

where 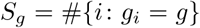, and 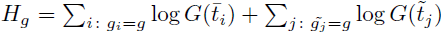. Clearly, the function 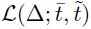 is differentiable in the whole parameter space (0, *∞*)^8^ and for each *g ∈ {*0, 1*}*^3^ the limits

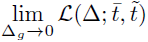

And

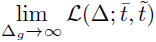

are both equal to *-∞*. Therefore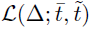 is maximized in a point where the partial derivatives 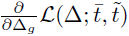 vanish. Fortunately

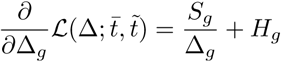

has unique zero at 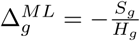, which gives the formulae for the maximum likelihood estimators. Fix the genotypes *g*_0_ *< g*_1_ *< g*_2_ *< g*_3_ and consider the set

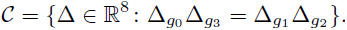

For the clarity of notation, as in the main text, we will denote the parameters of the null model in the set C as Δ^*C*^. Recall that the likelihood ratio test statistic is of the form

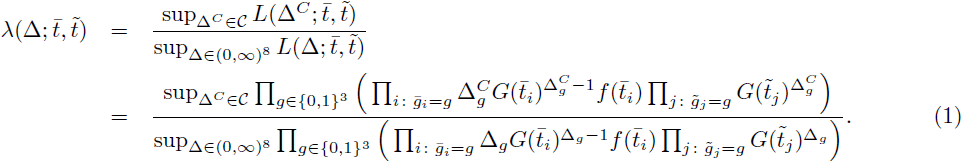

Clearly, the denominator is maximized for 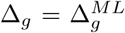. For the nominator, notice that for fixed 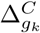 *k* = 0, 1, 2, 3 the likelihood 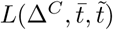 is maximal if the remaining 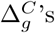 are the maximum likelihood estimators. It is therefore enough to maximize

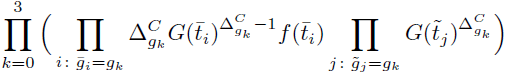

Over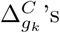 satisfying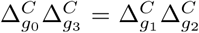. Equivalently we can maximize its logarithm, and of course w can omit constants in our optimization problem, which becomes

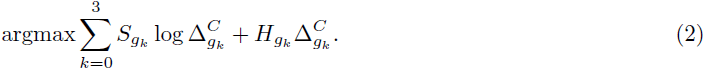

Let us replace sub-indices *g*_k_ by *k*, as in the main text. Substituting 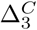 and 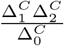 in (2) leads to

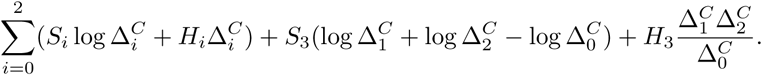

Let 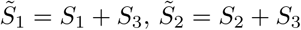 and 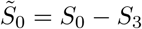. Our goal is to maximize the function

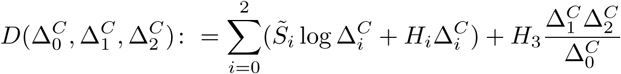

in Ω =(0, *∞*)^3^. Notice that lim_Δ*→∂Ω*_ *D*(Δ^*C*^) =-∞ and that *D* is differentiable on Ω. Thus it is enough to find zeros of its partial derivatives. Differentiating with respect to 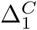 we obtain

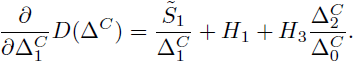

Comparing it to 0 we infer that

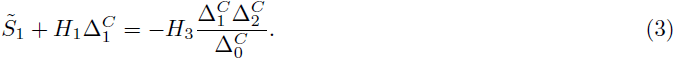

Similarly

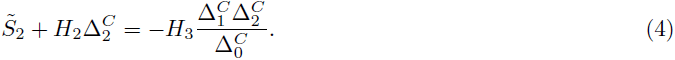

Equations (3) and (4) have the same right hand sides, hence

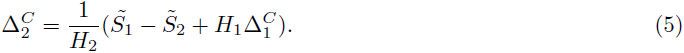

Applying the equality (5) into (3) leads to

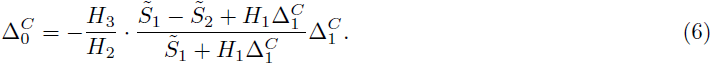

On the other hand, differentiating with respect to 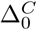 gives

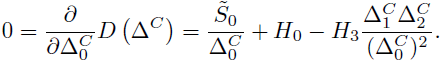

Multiplying by 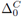 and substituting 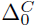 and 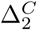 by the expressions from equalities (6) and (5) we obtain

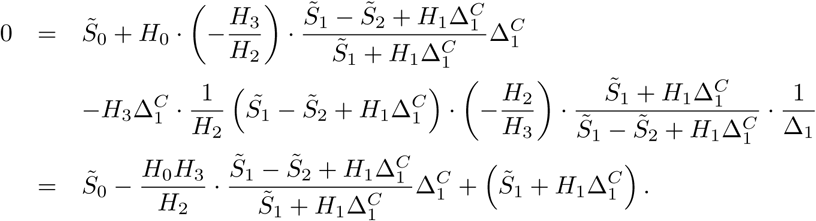

Multiplying by the denominators gives the following equivalent form

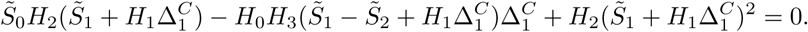

And after regrouping:

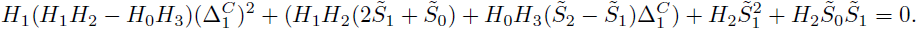

This equation, combined with equations (5), (6), and the condition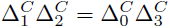, may have two solutions,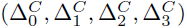 and 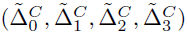, and we chose the one which lays in (0;∞)^4^. If both do, we choose the one for which the value of the expression

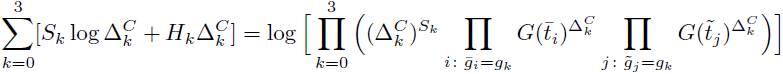

is greater.

Finally, inserting the estimated maximizing parameters into the nominator of the likelihood ratio (1), we obtain

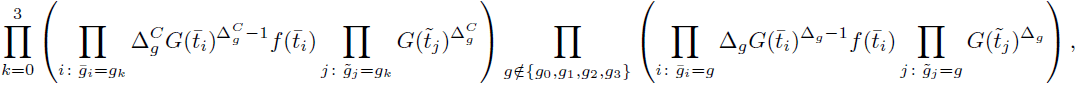

while inserting the estimated maximizing parameters into the denominator gives

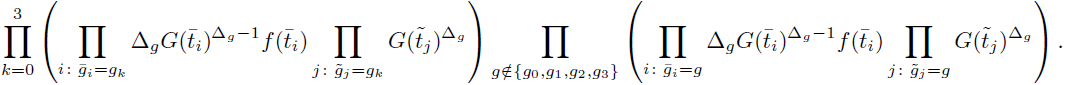

Thus, in the ratio (1) the expressions for g ∉ {g0, g1, g2, g3}, as well as the terms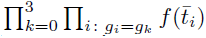 occur both in the nominator and in the denominator and cancel out. Multiplying both the nominator and the denominator by 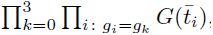, we obtain the likelihood ratio of the form presented in the main text

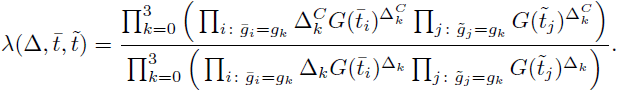

